# Development of a General Purpose Targeted LC-MS Method for Accurate Quantification of the SARS-CoV-2 Spike Protein Expression

**DOI:** 10.1101/2025.03.11.642564

**Authors:** Ruben Almey, Kevin Mwangi, Siegliende de Cae, Pathmanaban Ramasamy, Marijn Van Hulle, Ine Lentacker, Bert Schepens, Dieter Deforce, Geert Martens, Pieter Ramaut, Johannes PC Vissers, Xavier Saelens, Rein Verbeke, Maarten Dhaenens, Bart Van Puyvelde

## Abstract

The COVID-19 pandemic has catalyzed interest in immuno-multiple reaction monitoring (immuno-MRM) methods, with the detection of peptides unique to the nucleocapsid protein in nasopharyngeal swabs. While current applications predominantly focus on disease biomarkers, the pandemic has unveiled new opportunities, namely for the quantification of antigen expression following mRNA vaccination. Here, we present an optimized immuno-MRM method for quantifying SARS-CoV-2 spike protein fusion peptide, SFIEDLLFNK, for several practical applications. The method is versatile, applicable to multiple biological matrices, including plasma, and can be extended to nasopharyngeal swabs. It also offers a high-precision tool for assessing protein expression following plasmid and mRNA transfection. Moreover, in parallel to enabling accurate antigen quantification, the flow-through can be used to determine the proteome profile of the infected cells, providing insights into the intracellular immune response. This dual capability supports the rapid optimization of mRNA vaccines, thereby driving advancements in vaccine development strategies.

## Introduction

In the evolving landscape of SARS-CoV-2 vaccine development, mRNA-based vaccines have emerged as a transformative strategy, harnessing the host’s cellular machinery to generate a potent immune response against the virus ^1^. This approach offers distinct advantages, such as accelerated development timelines and the ability to be produced in cell-free systems. Currently, two main classes of mRNA vaccines are being developed: non-replicating mRNA and self-amplifying mRNA, each offering unique methods for antigen expression and vaccine delivery ^2^.

Although the concept of mRNA vaccines has been explored for over two decades, challenges related to stability, vulnerability to ribonuclease degradation, and efficient intracellular delivery have historically hindered progress ^3^. Key to advancing this technology are two critical factors: accurate mRNA concentration determination and lipid nanoparticle (LNP) design optimization. The latter plays a crucial role in protecting mRNA from degradation, enhancing cellular uptake, while accurate concentration determination ensures that the delivered dose is sufficient to provoke a strong and lasting immune response ^4^. Accurately quantifying spike protein expression, a key marker for vaccine potency, is therefore essential, as mRNA levels do not always directly correlate with protein abundance ^5,6^.

Traditional protein quantification techniques, such as Western blots and ELISA, have served as cornerstones in biomedical research. However, each method presents inherent limitations, ranging from gel-to-gel variations to batch-to-batch variability in antibody-based assays, compromising quantitative accuracy. In essence, the main concern is that antibodies are generated against the protein of interest and used to detect the target protein in non-native conditions in a complex protein mixture, giving rise to many sources of interference. Most of these disappear however when the immunogen and biomarker target become a simple linear tryptic peptide, which is how immuno-based strategies are combined with liquid chromatography coupled to mass spectrometry (LC-MS), almost entirely abrogating loss of binding and enabling protein detection within complex biological matrices ^7^. The synergy between peptide immuno-based techniques and MS do not only complement each other’s strengths, but also unlocks the potential for rapid and reliable quantification of protein targets ^8^.

An example of this integration is immuno-MRM, which was first commercially introduced by Stable Isotope Standards and Capture by Anti-Peptide Antibodies (SISCAPA) ^9^. In this approach, anti-peptide antibodies are immobilized onto magnetic beads, facilitating the capture of target peptide antigens. The incorporation of stable isotope labeled peptides (SIL) or a Quantification conCATamer (QconCAT) protein standard further refines the precision of protein measurement ^10^. For example, the use of this technology has been successfully established in several Clinical Laboratory Improvement Amendments (CLIA) compliant laboratories for thyroglobulin determination, aiding in the follow-up and monitoring of residual or recurrent disease in patients with differentiated thyroid cancer ^11–13^. Furthermore, it was applied by several groups for SARS-CoV-2 quantification, as an orthogonal approach to reverse transcription-quantitative polymerase chain reaction (RT-qPCR), to screen nasopharyngeal swabs for the presence of peptides unique to the nucleocapsid protein (NCAP) ^14–17^.

Here, we describe the continued development and validation of a high-throughput immuno-MRM method for the detection and quantification of a single peptide, i.e., SFIEDLLFNK, that acts as a surrogate for SARS-CoV-2 SPIKE protein. The application of the assay will be demonstrated for SARS-CoV-2 SPIKE quantitation in distinct matrices (plasma and nasopharyngeal UTM swabs) and reviewed in the context of plasmid and mRNA transfection efficiency measurement. Elegantly and separately presented, the analysis of antigen capture flowthrough enables rapid and comprehensive mapping of the mode of action and potential adverse outcome pathways, providing valuable insights to guide the future optimization of vaccine development.

## RESULTS

### A) Method development and validation

The development of an LC-MS-based assay for the SARS-CoV-2 spike (S) protein was initiated in 2020 by the CovMS consortium, a collaborative effort between academic and industrial research partners. Since then, multiple research groups have analyzed recombinant S-protein and published LC-MS methodologies, targeting tryptic peptides with the highest detectability (Supplementary Table 1)^18–33^. FLPFQQFGR and SFIEDLLFNK emerged as the most frequently detected. Interestingly, the CovMS consortium initially did not report FLPFQQFGR, potentially due to the omission of reduction and alkylation—a strategy originally employed to increase throughput. To address this, we repeated the digestion of pure recombinant S protein with reduction and alkylation, resulting in the detection of nearly all peptides listed in Supplementary Table 1, including FLPFQQFGR.

To date, several mutations have been identified in both peptide sequences and are catalogued in the Global Initiative on Sharing All Influenza Data (GISAID) EpiCov™ database (**Figure 1A**). However, none of these mutations are currently associated with variants of concern (VoC) or interest (VoI) ^34^. Notably, peptide SFIEDLLFNK was consistently detectable, regardless of reduction-alkylation, and exhibited superior MS detectability in-house compared to other S protein-derived peptides, making it the ideal candidate for targeted immuno-MRM (**Figure 1B**).

**Figure 1.**
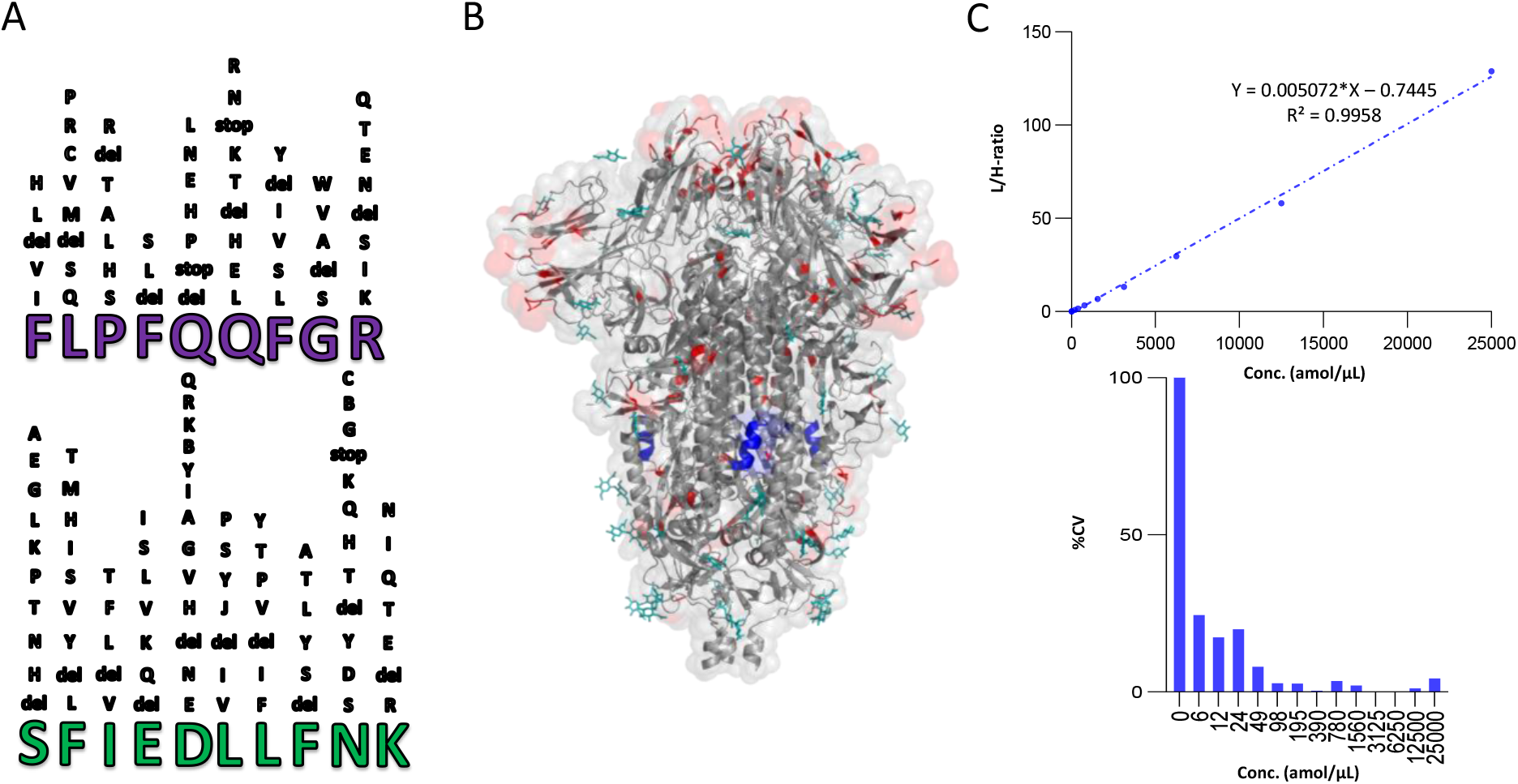
Selection and characterization of the SPIKE_SARS2 biomarker peptide. A) Combined analysis of publicly available and in-house data identified two consistently and reproducibly detected SPIKE_SARS2 peptides: FLPFQQFGR and SFIEDLLFNK. Mutations occurring in at least ten sequence entries from the GISAID database were mapped onto the peptide sequences, with the most frequent mutations displayed closest to their native amino acid positions. B) 3D structure of the SARS-CoV-2 SPIKE protein, highlighting the SFIEDLLFNK sequence (blue). All available natural variants (red) and glycosylation sites (turquoise) of the Spike glycoprotein (UniProt ID: P0DTC2), were retrieved from UniProt ^37^ and mapped onto the three-dimensional molecular structure (PDB ID: 6VXX) obtained from the RCSB Protein Data Bank ^38^. C) LLOD determination for peptide SFIEDLLFNK using a dilution series of synthetic endogenous SFIEDLLFNK peptide spiked into tryptically digested heavy-labeled QconCAT protein (3.5fmol/µL). A linear correlation (R^2^ > 0.99) between the Light/Heavy (L/H)-ratio and the peptide concentration was achieved. The percent coefficient of variation (%CV) values were calculated based on triplicate measurements obtained at each concentration.

For MS-based assays to move into a CLIA compliant environment, SIL internal standards are essential to meet the requirements for measurement accuracy ^35^. Here, a QConCAT, an artificial protein comprising concatenated peptides, already designed by the CovMS consortium in collaboration with PolyQuant, was used ^10^. Using this artificial protein allows for absolute quantification and enables performance evaluation between different laboratories. The ^15^N-labeled CovMS QconCAT protein, expressed in *E. coli*, incorporates an additional S protein biomarker peptide, GWIFGTLLDSK (AA 103-113), along with 11 nucleocapsid (NCAP) peptides, 3 RePLiCal peptides for optional retention time calibration, and 4 histone peptides ^18,36^. The inclusion of histone peptides allows for verification of proper nasopharyngeal swab application. Three fragment ions for the light peptide and two for the heavy QconCAT counterpart were selected and optimized for cone and collision energy to maximize sensitivity on a Xevo™ TQ-XS triple quadrupole mass spectrometer. Fragment ion selection prioritized achieving the highest signal-to-noise ratio to ensure robust and reliable quantification.

Performance was validated using a dilution series of the synthetic light peptide (0–25,000 amol/µL) spiked into digested CovMS QconCAT heavy standard (3.5 fmol/µL), measured in triplicate with a 2-min LC-MRM-MS method (**Figure 1C**). Peptide retention times demonstrated exceptional precision, with a standard deviation below 0.12 seconds. Good linearity (R^2^ = 0.9958) was observed even at low attomole concentrations, with light-to-heavy (L/H) peak area ratio reproducibility ranging from 0.25% to 24.70% coefficient of variation (%CV) (inset). The lower limit of quantification (LLOQ), defined as the lowest concentration where %CV was <20% and accuracy fell within 80–120%, was determined to be 390 amol/µL. Furthermore, the SFIEDLLFNK peptide remained detectable at concentrations as low as 6 amol/µL, aligning with previously reported LoD ^18^.

#### a. Nasopharyngeal swabs (patient samples)/Plasma

Following sensitivity validation of the LC-MS method using a synthetic peptide analogue in simple elution buffer (0.5% FA in water, 0.03% CHAPS), we incorporated an affinity-purified anti-SFIEDLLFNK polyclonal rabbit antibody to optimize sample preparation in two distinct matrices: nasopharyngeal swabs submerged in universal transport medium (UTM) and human plasma. The use of SISCAPA polyclonal antibodies conjugated to magnetic bead immuno-adsorbents enabled target peptide purification through a streamlined, addition-only protocol ideal for automation. This approach significantly enhanced throughput and minimized variability. An automated digestion and magnetic bead purification workflow was established using an Andrew+™ liquid handling robot (Waters Corporation, Milford, MA, USA), with a detailed deck layout provided in Supplementary Figure 1.

The best established immuno-MRM assay in-house (Cov^2^MS) was developed for diagnostic reasons and aimed to detect NCAP peptides in nasopharyngeal swabs and plasma ^15^. Therefore, we first made a back-to-back comparison with the existing workflow to assess the applicability of the new SPIKE target for this purpose. A recombinant S protein dilution series (0–50,000 amol/µL) was prepared in 180 µL of confirmed SARS-CoV-2 negative patient UTM (**Figure 2A**). The adapted protocol, included reduction and alkylation to improve tryptic digestion efficiency, which in turn could be assessed through the QconCAT internal standard ^15^. Although this addition extended sample processing time to 5 hrs and 36 min, it improved detection limits by approximately 20% (*data not shown*), likely due to increased digestion efficiency. Next, a single replicate peptide-only dilution series was prepared in plasma using the synthetic SFIEDLLFNK light peptide instead of a SPIKE recombinant protein (**Figure 2B**). An optimized version of the thyroglobulin SISCAPA plasma protocol was employed to increase the digestion efficiency of the plasma proteome (Material and Methods) ^11^. Both dilution series demonstrated excellent linearity (R^2^ > 0.99) down to 1 fmol/µL, closely matching the LLOQ established for the synthetic peptide in the simple elution buffer. Hence, the immuno-enrichment strategy effectively mitigated matrix effects, enabling reliable detection of the S protein in nasopharyngeal swabs and plasma.

**Figure 2.**
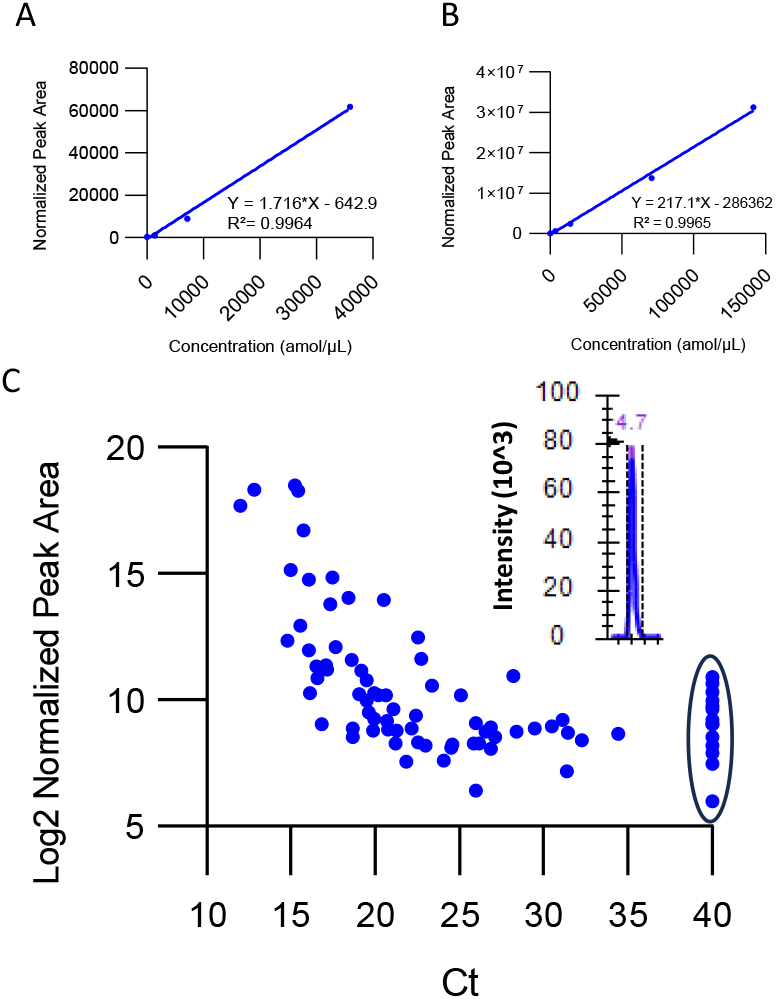
Performance of SISCAPA polyclonal anti-peptide antibody in Nasopharyngeal swabs (NP) and Plasma. A) A dilution series of recombinant S protein (0-36 fmol/µL) in UTM medium demonstrated excellent linearity with the normalized SFIEDLLFNK peak area (R^2^ = 0.9964). B) A dilution series of the synthetic SFIEDLLFNK peptide in plasma (0-140 fmol/µL) also showed strong linearity across a broad concentration range. C) Analysis of a cohort of 87 clinical nasopharyngeal swabs, where log2 normalized peak areas were compared to Ct values obtained via RT-qPCR. The MRM chromatogram for SFIEDLLFNK of a representative positive NP sample is shown inset. For simplicity, all RT-qPCR negative samples were assigned a Ct value of 40 (circled).

To assess its applicability in the analysis of NP swabs, a cohort of 87 nasopharyngeal swabs in UTM medium—73 classified as positive and 14 as negative by RT-qPCR—was processed and analyzed. The log2 normalized peak area for the SFIEDLLFNK peptide was plotted against the clinical cycle threshold (Ct) values obtained for the E-gene, as shown in **Figure 2C**. These two orthogonal analytical metrics were found earlier to correlate ^15^. At high viral loads (Ct < 20), a linear correlation was indeed observed between the output of both methodologies, consistent with findings previously reported for NCAP peptides. However, the linear correlation deteriorated at Ct values exceeding 20, as positive samples became indistinguishable from negative patient samples—indicated by the circled dots at Ct 40. Therefore, compared to the Cov^2^MS NCAP assay (Ct ~30-32), the S protein assay exhibited a 1000-fold lower detection limit ^15^. This disparity likely stems from the stoichiometric difference in viral particle encapsulation, where NCAP is considerably more abundant than the S protein ^39^. Interestingly, significant discrepancy in log2 normalized peak areas was also observed between samples with identical Ct values. While this could indicate biological differences—potentially correlating with symptomatology or disease severity—it is more likely attributed to repeated freeze-thaw cycles compromising sample integrity. An exemplary MRM chromatogram for SFIEDLLFNK from a highly positive NP patient sample (Ct 15) is depicted in the inset. The SFIEDLLFNK peptide represents a quantitatively accurate but less sensitive biomarker for SARS-CoV-2 detection in nasopharyngeal swabs compared to NCAP peptides used in the Cov^2^MS assay. Nevertheless, the immuno-enrichment strategy effectively enhances assay robustness and applicability, supporting its utility for S-protein analysis in clinical and research settings ^15^.

#### b. Quantifying recombinant SPIKE expression in HEK cells

Unlike viral detection, where stoichiometric differences between proteins pose challenges, mRNA vaccine-induced protein expression offers a more controlled and attractive application for the immuno-MRM assay targeting the SPIKE protein. As proof of concept, we evaluated the immuno-MRM assay against SFIEDLLNFK for monitoring recombinant S-protein expression in human embryonic kidney (HEK) cells, a commonly used system for protein expression due to its ability to generate essential post-translational modifications for proper protein folding and functional integrity ^40^. Here, a plasmid encoding the gene for the full-length spike protein was transfected with lipofectamine into HEK cells and two days after transfection, the cells were washed, snap-frozen, and processed for LC-MS analysis. The protocol described by Sutton et al. ^21^ was adapted to enhance compatibility with peptide immuno-enrichment through replacing Pierce™ RIPA Lysis buffer (containing SDS and SDC) with RIPA IP buffer.

Six biological replicates of SPIKE-transfected and empty plasmid transfected cell pellets were processed using our optimized immuno-enrichment protocol. Samples were analyzed via parallel reaction monitoring (PRM) on a TripleTOF 6600+ system. High S protein expression levels were observed in transfected cells (**Figure 3A, left**), with a normalized peak area reproducibility of 46.3 %CV. Notably, trace amounts of the S protein (1/4000) were detected in non-transfected control cell pellets (**Figure 3A, right**), likely due to minimal carry-over from previously analyzed transfected samples.

**Figure 3.**
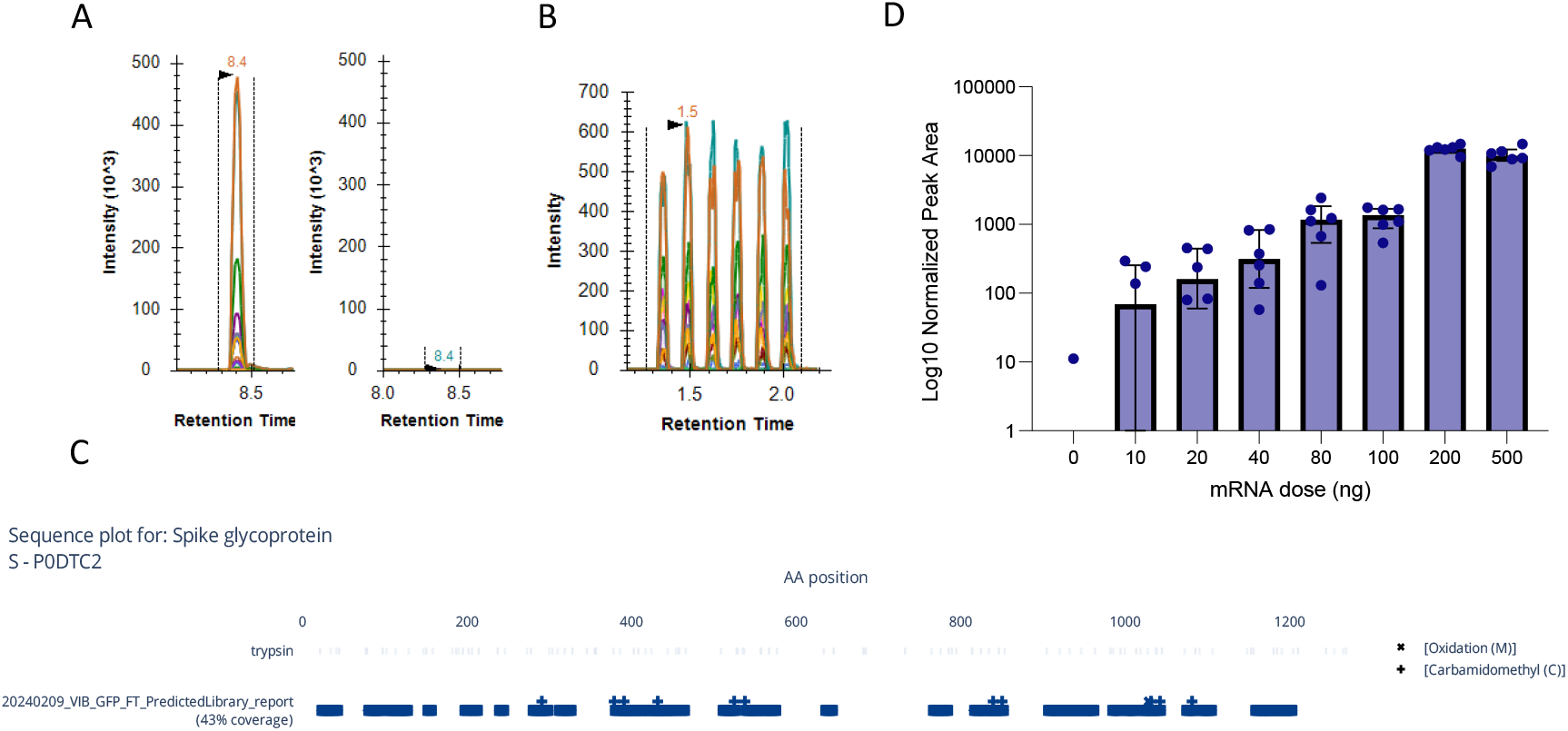
Transfection experiments with the HEK 293T cell line. A) HEK cells transfected with a SPIKE expression plasmid underwent SFIEDLLFNK peptide immuno-enrichment and were analyzed using LC-PRM (left panel). As a negative control, HEK cells transfected with an empty plasmid were analyzed (right panel) to ensure quality control. B) The positive samples were pooled and measured six times in PRM using AEMS. C) The flow-through obtained after antibody-bead incubation was subjected to solid-phase extraction (SPE) and analyzed in SWATH mode, covering 43% of the S protein sequence, figure generated with AlphaMap ^44^. D) A dose-response experiment was conducted by transfecting 30,000 HEK293T cells with varying amounts of the Comirnaty COVID-19 mRNA-LNP vaccine (0–500 ng). Following transfection, cells were analyzed for the presence of the SARS-CoV-2 spike protein peptide SFIEDLLFNK using LC-MRM. The bar graph shows the median log10 normalized peak area with interquartile ranges, based on six replicates per concentration.

Given the high expression levels observed, we assessed the potential for increasing throughput using acoustic ejection mass spectrometry (AEMS), which enables analysis of up to 2000 samples per hour with similar accuracy in quantification, only at the cost of sensitivity ^41^. Samples were analyzed using the Echo® MS+ system, which combines AEMS with a ZenoTOF 7600 MS system, achieving rapid sample readouts of 3.6 seconds per sample. Each sample was ejected six times and measured in PRM mode with 3-second intervals, resulting in highly reproducible total area measurements (%CV = 6%) (**Figure 3B**). However, the heavy-labeled peptide standard was undetectable due to its lower concentration relative to the endogenous peptide. Additionally, the protocol was not yet optimized for AEMS analysis: phosphate-buffered saline (PBS) was used as the washing buffer, and samples were eluted in 0.03% CHAPS with 0.5% formic acid, both of which suppress ionization and reduce AEMS sensitivity, as previously demonstrated 41.

MS is a versatile analytical tool and at this stage only the target peptide was measured. Yet, to mine the bulk of the available information in the sample, we also collected the flow-through (FT) after the 1-hour anti-peptide antibody incubation to screen for (i) additional S-protein peptides and (ii) the host response following transfection, on a high resolution QTOF instrument. The FT underwent solid-phase extraction (SPE) and was analyzed using data-independent acquisition (DIA) with the SWATH-MS approach on a TripleTOF 6600+ instrument. Peptide signals were extracted using DIA-NN software, employing a library-free method where a compiled FASTA database, containing Swiss-Prot reviewed SPIKE_SARS2 and human protein sequences, was *in silico* digested into tryptic peptides, and their retention times and fragment ion spectra predicted ^42^. This approach eliminates the need for spectral libraries generated through stochastic data-dependent acquisition (DDA), thereby enabling unbiased and comprehensive peptide and proteome coverage ^43^. A total of 34,813 peptide sequences were detected, of which 52 peptides mapped to the S protein, covering 43.1% of the wild strain protein sequence (**Figure 3C**). Interestingly, the SFIEDLLFNK peptide was still detectable in the FT after immuno-enrichment, likely due to saturation of the antibody epitope binding sites, consistent with the high cell count (~1 million) used for S protein expression.

In addition to the S-protein peptides, the remaining 34,761 peptide sequences matched 5,564 human protein sequences. Importantly, when the purified peptide was measured by untargeted DDA acquisition, only 1 peptide could confidently be annotated (*data not shown*). This illustrates that the full proteome sample without the immunopurified target peptide is still present in the FT. Therefore, an untargeted discovery proteome analysis can be conducted to obtain valuable information about the mode of action or adverse outcome pathways. Unfortunately, the initial experiment was not designed for this and did not allow for robust statistical interpretation of changes that are induced in the host cell upon transfection and is therefore only reported to illustrate proteome detectability purposes.

#### c. mRNA expression in HEK cells

After confirming the successful measurement of recombinant S protein expression in plasmid-transfected HEK cells using the immuno-MRM assay, we proceeded to assess protein expression following transfection with mRNA lipid nanoparticles (LNPs). Compared to plasmid-based expression vectors, mRNA transfection bypasses the need to cross the nuclear barrier, allowing for more immediate protein expression. However, the duration of expression is shorter due to the transient nature of mRNA, which is more rapidly degraded. Therefore, we conducted a dose-response experiment (0–500 ng) using the Pfizer-BioNTech COVID-19 Omicron XBB.1.5 mRNA vaccine (Comirnaty, New York, USA) on 30,000 HEK293T cells per well. The mRNA dosage was converted from nanograms to femtomoles to allow direct comparison between the amount of mRNA transfected and the spike protein expressed, as suggested by Sutton et al ^21^. This conversion yielded mRNA transfection doses ranging from 7.1 fmol to 357.1 fmol. The mRNA concentration was validated using a RiboGreen assay to ensure accuracy.

After 24 hrs of incubation under optimal growth conditions, cells were harvested by centrifugation, supernatant was removed, and the resulting pellets were snap-frozen. The optimized sample preparation protocol previously used for plasmid-transfected HEK cells was applied. Peptides obtained after immuno-enrichment were separated using a 2-minute LC gradient and quantified in MRM mode on the Xevo TQ-XS instrument. The resulting chromatograms for the different dosages are provided in Supplementary Figure 2. To confirm effective sample clean-up, two peptides from the nucleocapsid (NCAP) protein— ADETQALPQR and AYNVTQAFGR—were included as negative controls. As expected, no measurable signal was detected for these peptides in their light and heavy-labeled form.

As shown in **Figure 3D**, measurable S protein expression was observed even at the lowest mRNA LNP dose (10 ng), though the signal was below LLOQ, with a %CV of 131.7%, reflecting the inherent challenges of low expression levels with mRNA transfection. Increasing mRNA concentrations led to higher protein expression levels and improved reproducibility, as indicated by reduced %CV values. Notably, transfection with 200 ng (142.9 fmol) of mRNA resulted in the highest protein expression levels and excellent biological reproducibility, with a %CV of only 13.9%. Using the SFIEDLLFNK peptide dilution series in elution buffer (**Figure 1C**), we estimated that transfection with 200 ng of mRNA yielded a spike protein concentration of 24-49 amol/µL in 50 µL elution buffer, corresponding to a total expressed S protein amount of approximately 1.2 to 2. 5 fmol. This equates to a transfection efficiency of roughly 1% of the mRNA after 24 hrs of incubation. Notably, protein expression declined at higher mRNA doses (500 ng), with a similar trend observed at 1000 ng (*data not shown*), likely due to saturation of the translational machinery or cellular stress responses induced by elevated mRNA levels. To further study this, the workflows has the option to assess the impact on the host cell by measuring the flow-through of proteins after iMRM. Taken together, absolute quantification of protein expression after mRNA transfection with LNPs can be very informative during the therapeutic development process.

#### d. Assessing the impact on the host cell

To investigate the mode of action and adverse outcome pathways, we analyzed the flow-through (FT) fractions from the NTC (non-transfected control), 200 ng, and 1000 ng transfection experiments using ZenoSWATH acquisition on a ZenoTOF 7600 system. This workflow reproducibly identified over 4000 proteins in the FT fraction. Principal Component Analysis (Supplementary Figure 3A) confirmed clustering of samples by condition, with the exception of one outlier in the 200 ng group, indicating overall consistency in the data. Additionally, CV analysis (Supplementary Figure 3B) showed that peptide and protein variability remained below 20% across all conditions, further supporting the reproducibility and reliability of the data. Subsequent GO enrichment analysis of differentially expressed proteins revealed a downregulation of genes associated with cytoplasmic translation in transfected cells compared to the NTC samples (Supplementary Figure 3C). Tatematsu et al. already reported that exogenously delivered mRNA is intrinsically immunogenic, triggering several innate immune sensing pathways, which leads to the production of inflammatory cytokines and suppression of cellular translation, undesirable for the production of the therapeutic protein ^45^. Although modified nucleotides have been used in the COVID-vaccines to mitigate innate immune activation, some residual triggering persists ^46^. Furthermore, we noted that the cells transfected with 1000 ng revealed significant downregulation of key RNA-binding and chaperone proteins, including PA2G4, HSP90AB1, HNRNPU, KHSRP, MATR3, RBMX, NONO, and RBM14. These proteins are essential for mRNA processing, stabilization, and translation, suggesting that their depletion may impair the cellular capacity to efficiently translate exogenous mRNA at high concentrations. This observation aligns with findings from the SARS-CoV-2 RNA interactome study, which demonstrated that viral RNA hijacks similar host proteins to facilitate its replication while disrupting normal RNA homeostasis ^47^. The overlap between the interactome and our data highlights a possible shared mechanism, where stress-induced downregulation of RNA-binding proteins leads to sequestration or degradation of LNP mRNA, ultimately limiting translation. These results underscore the delicate balance of host cellular pathways required for optimal mRNA translation and provide insights into the saturation effects observed at higher mRNA doses.

This intracellular immune response can take many forms and is dependent on the transfection agent used. To further explore host cell responses, we conducted a more complex experiment including two transfections with the commercial transfection agent lipofectamine comparing the delivery of nucleoside-modified mRNA with two distinct self-amplifying RNA (saRNA) constructs and measured the full proteome as described above. Importantly, SPIKE measurement was not performed in this experiment, as the constructs used did not encode the respective sequences. The sole purpose of this additional experiment was to assess intracellular immune responses under different transfection conditions. We compared mock transfected and HeLa cells transfected with conventional modified mRNA and two different saRNA constructs in six replicates each. The detailed report of the analysis that was created with MS-DAP can be found as supplementary data ^48,49^. Briefly, close to 5000 proteins were identified in each sample, with over 50,000 peptides identified in over 25% of the runs and median %CVs below 15%. The comparison between mock transfected, nucleoside-modified mRNA transfected and two different saRNA constructs showed clear separation in the PCA (**Figure 4A**). Indeed, the most prominently upregulated proteins in the saRNA transfected cells were STAT1, activating the transcription of IFN-stimulated genes (ISG), and ISG15 itself, which drives the cell in an antiviral state by tagging proteins for degradation in viral-infected cells (**Figure 4B** and Supplementary Figure 4). In turn, these changes can be indications of the presence of double stranded RNA, which could steer the future development process of therapeutics that are based on this technology.

**Figure 4.**
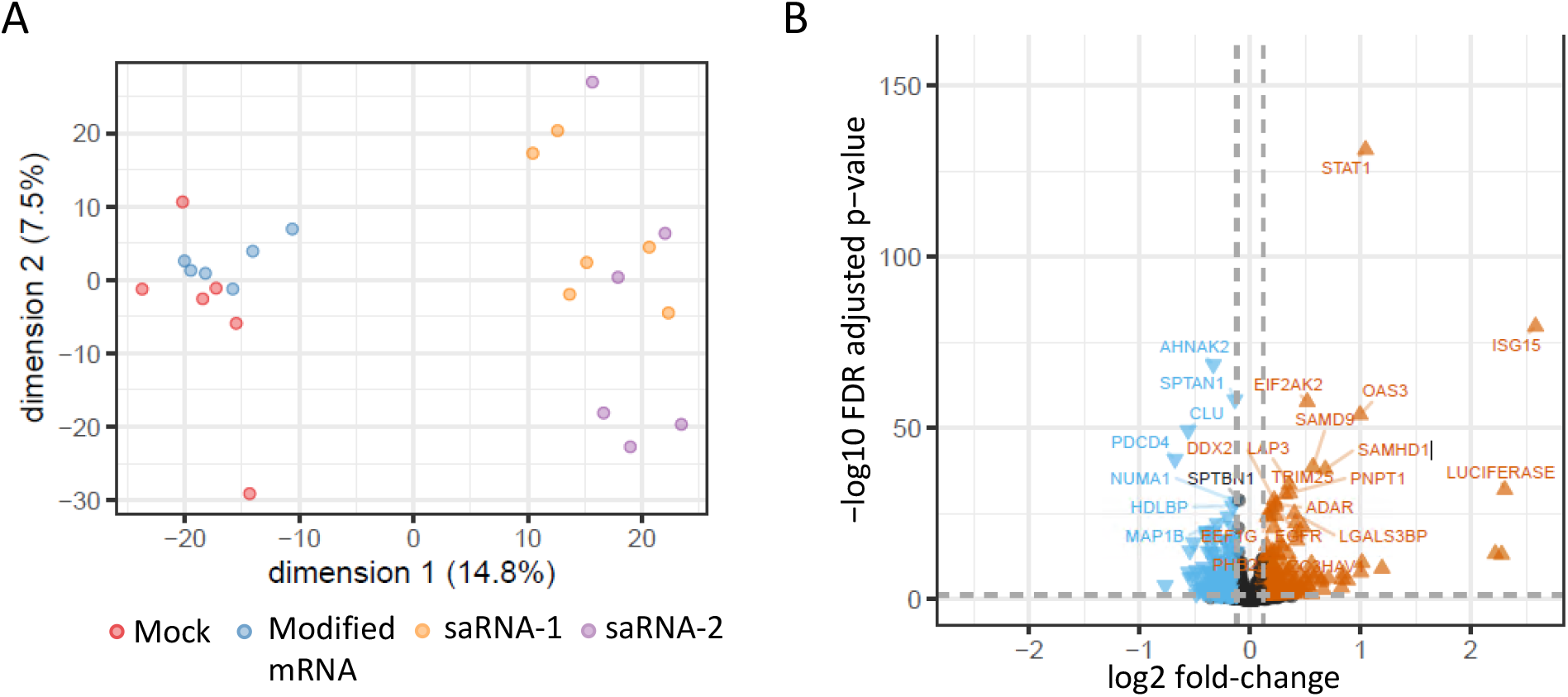
Lipofectamine transfection experiment with Mock, Modified mRNA and two different saRNA constructs. A) Principal Component Analysis of the untargeted proteomics dataset, showing primary sample clustering based on experimental conditions: Mock transfection (red), Modified mRNA transfection (blue), and two different saRNA constructs: saRNA-1 (orange) and saRNA-2 (purple). Percent variance explained by each principal component is indicated on the axes. Clear separation of clusters suggests distinct proteomic profiles for the conditions. B) Volcano distribution highlighting differential protein expression, using MSqRob, between mRNA and saRNA transfected cells. The x-axis represents the log2 fold change, and the y-axis represents −log10 FDR adjusted p-value. The 25 most significantly upregulated (orange) and downregulated (blue) proteins in the saRNA group are annotated, with STAT1_HUMAN and ISG15_HUMAN prominently upregulated in the saRNA group. Non-significant abundance changing proteins are shown in black.

In conclusion, combining targeted immuno-MRM for S protein expression measurement with untargeted DIA analysis of the FT, provides a comprehensive assessment of both transfection efficiency and the resulting cellular immune response. This approach holds promise for monitoring immune activation following mRNA/saRNA vaccination, offering a deeper understanding of antigen behavior within host cells. These insights are critical for evaluating RNA-based therapies and optimizing their performance in preclinical and clinical studies.

## DISCUSSION

One of the most significant advancements catalyzed by the COVID-19 pandemic has been the rapid introduction of mRNA vaccines into clinical practice. These vaccines have revolutionized the field of vaccinology by harnessing the host’s cellular machinery to produce viral proteins, such as the SARS-CoV-2 spike protein, to stimulate a protective immune response. The success of mRNA vaccine platforms for SARS-CoV-2 has underscored the importance of developing robust and precise methods for quantifying protein expression following vaccination, particularly as this technology will expand to address other infectious diseases in the near future—such as influenza and respiratory syncytial virus (RSV) ^50,51^.

In this study, an optimized immuno-MRM assay for the sensitive detection of the fusion peptide SFIEDLLFNK was employed, which served as a surrogate marker for SARS-CoV-2 spike protein expression. An excellent correlation between the concentration of pure synthetic peptide and the L/H-peak area was demonstrated, spanning four orders of dynamic range (R^2^ = 0.996). This high level of assay sensitivity enabled the detection of S-protein expression even at the lowest tested dosage, i.e. 10 ng of mRNA-LNP. While our method demonstrates robust performance in a cell-based model, further validation is required in *in vivo* settings to ensure its applicability for monitoring vaccine-induced protein expression in real-world scenarios. This is particularly important given that data on the biodistribution of mRNA and the resulting protein production in *in vivo* environments remains limited ^52^. Additionally, the method developed in this study could also be applied to assess vaccine stability under various storage and transportation conditions, including extreme temperature variations. Given the susceptibility of mRNA-LNP vaccines to degradation, the immuno-MRM method could serve as a potency assay to monitor protein integrity post-administration ^53^.

Furthermore, recent studies have underscored the importance of monitoring (circulating) protein levels following mRNA vaccination, particularly in the context of vaccine-induced adverse effects. Elevated spike protein concentrations have been observed in individuals with myocarditis following COVID-19 vaccination, and persistent S-protein presence has been linked to Long-COVID symptoms ^54,55^. These findings underscore the necessity of developing assays capable of detecting low levels of vaccine-induced proteins, as they can provide critical insights into the safety and long-term effects of mRNA vaccines. Although we demonstrated the detection of SFIEDLLFNK synthetic peptide in plasma, we were unable to detect distinct MRM signals in plasma samples from individuals vaccinated with a Covid-19 mRNA vaccine (*data not shown*). Despite our optimized enrichment strategy being applied to 200 µL of plasma, other studies have successfully identified spike protein biomarkers without enrichment using dried blood spots (DBS) ^56^. According to Cosentino et al. ^57^, circulating S-protein concentration 24 hrs post-mRNA vaccination are estimated to reach approximately 150 pg/mL (~1.5 fmol/mL), which, in our assay, would correspond to 6 amol/µL in a 50 µL elution buffer—below the limit of detection (LoD) of our assay^57^. Nonetheless, higher concentrations of circulating S-protein have been reported in individuals who experienced myocarditis following Covid-19 mRNA vaccination, as well as in patients with Long Covid.

Finally, very little is known about the immune response from the host cell upon infection. We show here that the flow through after immuno-enrichment of the target peptide can be measured using a fast (20 min) untargeted acquisition method like DIA to better understand the expressional changes induced in the host cell, as demonstrated for the saRNA transfection. Indeed, these seem to be mainly related to translation machinery and immune response to viral infection, both unwanted changes that can limit the amount of antigen presented to the immune system post-vaccination. This parallel measurement on the same sample will therefore prove of great value to predict and minimize unwanted side effects in future mRNA- or saRNA-based vaccines.

In conclusion, the ability to measure protein expression accurately after RNA-based vaccination is paramount, both for optimizing vaccine efficacy and for monitoring potential adverse effects. As mRNA vaccines for diseases beyond COVID-19 are developed, robust and sensitive assays like the one presented here will play an indispensable role in ensuring the success and safety of these next-generation vaccines.

## Material and Methods

### Reagents and materials

Unlabeled (light) and stable isotope labeled (SIL or heavy) peptides for SFIEDLLFNK were obtained from Biosynth (Staad, Switzerland). Upon receipt from the vendor, the lyophilized peptides were solubilized in 30% (ACN)/0.1% formic acid (FA) and stored frozen at −80°C. The CovMS QconCAT standard was kindly provided by Polyquant (Bad Abbach, Germany).

Polyclonal anti-peptide antibodies against SFIEDLLFNK were produced by SISCAPA Assay Technologies (Victoria, Canada) and were in-house coupled to magnetic Dynabeads Protein G (Thermo Fischer Scientific, MA, USA). Recombinant SPIKE_SARS2 protein (2019-nCov) was produced in insect cells with a baculovirus expression system (Sino Biological, Beijing, China). Tosyllysine Chloromethyl Ketone (TLCK), Dimethyl Pimelimidate dihydrochloride (DMP), Sodium azide bioultra, Tris(2-carboxyethyl)phosphine hydrochloride solution (TCEP) and 3-[(3-Cholamidopropyl)dimethylammonio]-1-propanesulfonate (CHAPS) were sourced from Sigma-Aldrich (St. Louis, MI, USA). Iodoacetamide (IAA) and Pierce™ RIPA IP Lysis buffer were obtained from Thermo Fischer Scientific (Waltham, MA, USA). Trypsin/Lys-C and TPCK treated Trypsin were sourced from Promega (Madison, Wisconsin, USA) and Worthington Biochemical (Lakewoord, NJ, USA).

Residual Covid-19 nasopharyngeal swab and plasma samples were obtained from the AZ Delta Hospital, Roeselare, and Ghent University Hospital, respectively, with approval of the University Hospital Ghent ethics committee (BC-09263 and BC-11711). This study was performed in accordance with the Helsinki declaration.

### Antibody bead coupling

First, the required amount of Protein G Dynabeads (30 mg/mL), namely 500 µL, were washed three times using a wash buffer composed of 1x PBS + 0.03% CHAPS. 100 µg of the polyclonal antibody mixture against the SFIEDLLFNK peptide (0.25 mg/mL) was transferred to a 15 mL falcon tube, after which the required amount of washed magnetic protein G beads is added. The antibody/bead mixture is further diluted with wash buffer to an antibody concentration of approximately 0.01 mg/mL, after which it is incubated at 4°C with end-over-end mixing. After overnight incubation, the tube is placed in a custom-made magnetic separation rack to pull the beads to the side, after which all the liquid is removed. 15 mL of a crosslinking solution, composed of 20 mM DMP in 200 mM triethanolamine (pH 8.5), was added followed by an end-over-end ambient temperature mixing step for 30 min. The tube was again placed on a magnet to pull the beads to the side of the tube after which all the volume is aspirated. 15 mL of a 150 mM monoethanolamine (pH 9) solution was added to quench the reaction. After a 30 min incubation at RT, again all the volume was removed with the tube on the magnet. Any residual unbound material was washed away using 15 mL of an acidic wash solution (0.03% CHAPS in 0.1% formic acid). Since this washing step could result in the uncoupling of the antibody, the tube was removed from its volume immediately after all beads were in solution through vortexing. To remove any acidic residue, the mixture was washed twice with wash buffer, after which the crosslinked antibodies were resuspended in 1xPBS (0.03% CHAPS, 0.01% NaN3) to a concentration of 0.1 µg/µL, followed by storing them at 4°C.

### Peptide enrichment

Prior to the addition of the antibody-coupled magnetic bead immuno-adsorbents, a step was included to fully resuspend the beads, after which 10 µL (1 µg) was added to the digested samples. After antibody-bead addition, washing and elution was performed as previously described ^58^. In short, the antibody-bead mixture was washed three times with 150 µL 0.03% CHAPS in 1x PBS, followed by elution in 50 µL of 0.03% CHAPS 0.5% FA in Water.

### Sample preparation

#### 1) Nasopharyngeal swabs

Residual Covid-19 nasopharyngeal samples were obtained from the AZ Delta Hospital, Roeselare, Belgium, with approval of the University Hospital Ghent ethics committee (BC-09263). The Nasopharyngeal swabs were prepared using a slightly altered protocol described by Van Puyvelde et al. (2022), wherein proteins in 180 µL of undiluted Bioer Universal Transport Medium (UTM) underwent protein precipitation by the addition of 1260 µL of ice-cold acetone in a 1.5 mL protein LoBind Eppendorf tube. After centrifugation (16,000g) in a cold environment (0°C) for 10 min, the supernatant was discarded and the residual protein pellet was resuspended in 100 µL of a 100 mM ABC solution, spiked with 250 fmol of the QconCAT CovMS protein which contains a heavy labelled version of the SFIEDLLFNK peptide. The tubes were extensively vortexed prior to transferring the volume to a 700 µL round-well collection plate (Waters Corporation). The next steps in the protocol were performed using an Andrew+ automation system to reduce variation introduced by manual pipetting. First, 5 µL of a 101 mM TCEP solution was added, followed by a 30-min incubation at 37°C. After cooling down the plate, 5 µL of a 303 mM IAA solution was added to alkylate the free cysteines during a 15-min incubation at room temperature (RT). 20 µL of a 0.05 µg/µL trypsin-Lys C solution (Promega, Madison, WI, USA) in 100 mM NH4HCO3 was added, followed by a 60-minute incubation at 37°C. Trypsin activity was inhibited by adding 20 µL of a 0.22 mg/mL TLCK solution in 10 mM HCl, followed by SFIEDLLFNK enrichment using a polyclonal antibody mixture (see peptide enrichment). A total of 87 nasopharyngeal patient samples, of which 73 were confirmed to be positive for SARS-CoV-2 using the STARMag Viral DNA/RNA 200 C RT-qPCR assay (Seegene Technologie, Walnut Creeck, CA, USA), were prepared using this protocol, alongside a dilution curve comprising 8 samples.

#### 2) Plasma

Remnant plasma samples from whole blood collected in anticoagulant-treated (EDTA) tubes, was obtained from healthy volunteers with approval of the University Hospital Ghent ethics committee (BC-11711). Since plasma is a more complex medium compared to nasopharyngeal swabs, a modified version of the SISCAPA thyroglobulin protocol was employed. Briefly, 200-400 µL of plasma or serum was spiked with 250 fmol of QconCAT protein. This mixture underwent a 40-minute denaturation and reduction process in 2% sodium deoxycholate (SDC) and 10 mM tris(2-carboxyethyl)phosphine (TCEP) in 0.2M TRIZMA at 37°C with agitation at 600 rpm. Alkylation was then performed using iodoacetamide at a final concentration of 10 mM, followed by a 40-minute incubation at room temperature. Subsequently, 50 µL of 10 µg/µL trypsin-TPCK (Worthington, New Jersey, USA) in 0.2 M TRIZMA was added, and the sample was incubated for 3 hrs at 37°C. Trypsin activity was inhibited by adding 20 µL of a 3 mg/mL TLCK solution in 10 mM HCl. Following a 30-minute incubation, enrichment of the SFIEDLLFNK peptide was performed as described earlier (see peptide enrichment).

#### 3) Transfection experiments

##### a. Plasmid transfection

HEK293-T cells (ATCC; CRL-3216) and Vero E6 cells (ATCC; CRL-1586) were cultured at 37°C in the presence of 5% CO2 in DMEM supplemented with 10% heat-inactivated FBS, 1% penicillin, 1% streptomycin, 2 mM L-glutamine, non-essential amino acids (Invitrogen) and 1 mM sodium pyruvate. One day before transfection HEK293-T cells were seeded in 6-well plates at 200 000 cells per well. Six wells of cells were each transfected with 750 ng of a SARS-CoV-2 spike(del-18) expression plasmid in combination with 250 ng of an GFP expression plasmid using 3 µL Fugene according to the manufacturers guidelines (Promega) ^59^. As controls six wells of cells were transfected with an empty expression vector. Two days after transfection the growth medium was removed and the cells detached by using 1 ml PBS. The cells were centrifuged and after removal of the supernatant the resulting 12 pellets were snap frozen and stored at −80 °C.

##### b. mRNA-LNP transfection | Potency Assay

HEK293-T cells (ATCC; CRL-3216) for transfections were grown in DMEM (Gibco) supplemented with 10% FBS and 5% penicillin and streptomycin (Gibco) at 37 °C and 5% CO2. All cells tested negative for mycoplasma presence using MycoAlert Mycoplasma Detection kit per the manufacturer’s specifications (Lonza). When the HEK293T cells reached approximately 80% confluency, cells were seeded into a flat bottomed 96-well plate at a density of 30,000 cells in 200 µL per well. After 24hrs of seeding at 37°C to allow for proper cell attachment, the cells were spiked with 0 ng, 10 ng, 20 ng, 40 ng, 80 ng, 100 ng, 200 ng and 500 ng of Pfizer Comirnaty™ mRNA vaccine (Omicron XBB.1.5) each having 6 biological replicates. After 24hrs at 37°C in a humidified atmosphere with 5% CO_2_, the culture medium was aspirated, and the cells were carefully washed with PBS to avoid cell detachment. Following the wash, 0.25% trypsin-EDTA was added to facilitate detachment. Trypsin activity was neutralized by adding fresh culture medium and finally the cells were transferred to a new U-bottom 96-well plate prior to freeze drying.

The protocol from Sutton et al. was optimized to make it compatible with peptide immuno-enrichment ^21^. In short, to each well of the plate 180 µL of Pierce™ RIPA IP Lysis buffer was added, followed by mixing repeatedly to detach the cells from the bottom of the well. After a 30-min shaking step, the cells were sonicated for 5 min using the PIXUL® ultrasonicator (50N pulses, 1 kHz frequency and 20 Hz burst rate) to obtain complete lysis of the cell pellet. Following sonication, the liquid was transferred to 1.5 mL protein LoBind tubes and 1260 µL of ice-cold acetone was added to precipitate the proteins. After thorough vortexing, samples were centrifuged at 16,000 g for 10 mins at 0°C, followed by removing the supernatant. The dry pellets were resuspended in 50 µL of 50 mM ABC buffer containing 0.1% RapiGest™ SF surfactant, followed by 207.6 fmol spike of the QconCAT CovMS standard. Proteins were reduced by incubating samples with 10 mM Tris(2-carboxyethyl)phosphine (TCEP) at 37°C for 30 min. Alkylation was subsequently performed by adding 10 mM iodoacetamide (IAA) and incubating the mixture at room temperature for 15 min, protected from light to prevent IAA degradation. Digestion was performed for 1 hr at 37°C using 1 µg of Trypsin/Lys-C, followed by a 5-min trypsin incubation step using 20 µL of a 0.22 mg/mL TLCK solution. Subsequent enrichment of the target peptide was carried out following the protocol detailed in the ‘Peptide Enrichment’ section.

##### c. saRNA transfection

saRNA constructs were based on the VEEV TC-83 genome with an A3G mutation in the 5’UTR. The structural proteins were replaced by a firefly luciferase coding sequence. saRNA-1 contains the wild type nsP sequence, while saRNA-2 contains an nsP2 sequence with the Q739L mutation, which was previously shown to cause less cytopathogenicity in transfected cells ^60^. Plasmids encoding the saRNAs were linearized by digestion with BspQI to generate an IVT template. The linearized plasmid was used as an IVT template with the HiScribe T7 High Yield RNA Synthesis Kit (NEB). Next, the template DNA was digested using Turbo DNAse for 30 minutes (ThermoFisher). The resulting IVT saRNA was then purified using the Monarch RNA Cleanup Kit (NEB).

HeLa cells (ATCC; CCL-2) were grown in MEM (Gibco) with 10% FBS, 1% penicillin and streptomycin, 1 mM Sodium Pyruvate and 1% MEM NEAA at 37°C and 5% CO2. 250 000 HeLa cells were seeded per well in 6 well plates. After 24 hours the cells were transfected with RNA using Lipofectamine MessengerMax (ThermoFisher) (250 ng/ well) at a RNA:lipofectamine ratio of 1:2. 6 replicates were included for each condition. At 24 hours post transfection the culture medium was aspirated, and the cells were carefully washed with PBS to avoid cell detachment. Following the wash, 0.25% trypsin-EDTA was added to facilitate detachment. Trypsin activity was neutralized by adding fresh culture medium and finally the cells were transferred to 1.5 ml tubes. Cell pellets were spun down and washed twice with ice cold PBS, after which the cell pellets were flash frozen in dry ice with ethanol.

### SPE

For several experiments, the flow through (FT) obtained after peptide enrichment was retained and cleaned-up by SPE for further analysis. For this, the FT after SISCAPA enrichment was acidified by adding several µL of 100% FA, after which it was loaded to 1 mL Strata-X 33 µm Polymeric Reversed Phase cartridges (Phenomenex). The cartridges were preconditioned with 100% methanol and equilibrated twice with milliQ water. After sample loading, the columns were washed twice with 5% MeOH in water, followed by two consecutive peptide elution steps in 60% acetonitrile (ACN). The obtained samples were vacuum dried in a SpeedVac and resuspended in 0.1% FA in water prior to LC-MS analysis.

**Table 1.** MRM parameters for target peptides.

| Peptide | Precursor m/z | Fragment m/z | Cone Voltage (V) | Collision Energy (V) |
| --- | --- | --- | --- | --- |
| ADETQALPQR (NCAP) | 564.78 | 400.23 | 20 | 17 |
|  | 564.78 | 584.35 | 20 | 16 |
|  | 564.78 | 712.41 | 20 | 17 |
|  | 572.26 | 407.21 | 20 | 17 |
|  | 572.26 | 593.32 | 20 | 16 |
| AYNVTQAFGR (NCAP) | 563.79 | 349.15 | 30 | 17 |
|  | 563.79 | 679.35 | 30 | 16 |
|  | 563.79 | 892.46 | 30 | 17 |
|  | 571.26 | 353.14 | 30 | 17 |
|  | 571.26 | 689.32 | 30 | 16 |
| SFIEDLLFNK (SPIKE) | 613.33 | 408.22 | 40 | 25 |
|  | 613.33 | 878.46 | 40 | 19 |
|  | 613.33 | 991.55 | 40 | 17 |
|  | 619.31 | 887.44 | 40 | 19 |
|  | 619.31 | 1001.52 | 40 | 17 |

### LC-MS analysis

#### MRM

MRM analyses were performed on an ACQUITY™ UPLC™ I-Class system coupled to a Xevo TQ-XS instrument (Waters Corporation, Wilmslow, UK). Ten microliter of each sample was injected after which peptide separation was performed using a gradient elution of mobile phase A containing LC–MS-grade deionized water with 0.1% (v/v) formic acid and mobile phase B containing LC–MS-grade acetonitrile with 0.1% (v/v) formic acid. The Xevo TQ-XS conditions were as follows: capillary voltage 0.5 kV, source temperature 120 °C, desolvation temperature 600 °C, cone gas flow 150 L/h, and desolvation gas flow 1000 L/h. The MS was calibrated at unit mass resolution for MS1 and MS2. Endogenous and the corresponding stable isotope labelled peptides were detected using MRM acquisition using the following experimental settings.

For the 8-min run, gradient elution was performed at 0.6 mL/min using an ACQUITY Peptide BEH C18 column (2.1 mm × 50 mm, 1.7 μm, 300 Å) with initial inlet conditions at 5 % B, increasing to 33 % B over 5.5 min, followed by a column wash at 90 % B for 1.4 min and a return to initial conditions at 5 % B.

For the 2-min run, gradient elution was performed at 0.8 mL/min using an ACQUITY Premier Peptide BEH C18 column (2.1 mm × 30 mm, 1.7 μm, 300 Å) with initial inlet conditions at 5 % B, increasing to 15 % B from 0.15 to 0.35 min and at a steady state for 0.25 min. Subsequently, over 0.4 min, gradient B was increased to 25%, followed by a column wash at 90 % B for 0.25 min and returning to initial conditions at 5 % B. The total run time was 1.8 min.

#### PRM

##### A) TripleTOF 6600+

An Eksigent NanoLC 425 System (SCIEX, Massachusetts, USA) plumbed for microflow chromatography was used and operated in trap-elute with the SCIEX TripleTOF 6600+ mass spectrometer. Peptides were trapped on a YMC TriArt C18 guard column (id 500 µm, length 5 mm, particle size 3 µm) and separated using a 150 × 0.3 mm, 2.6 µm, 100 Å Luna Omega Polar C18 Column (Phenomenex, California, USA), temperature controlled at 30 °C. Mobile phase A consisted of UPLC-grade water with 0.1% (v/v) FA, and mobile phase B consisted of UPLC-grade ACN with 0.1% (v/v) FA. Peptide elution was performed at 5 µL/min using the following gradient: i) 3% to 40% mobile phase B in 6 min, ii) ramp to 80% mobile phase B in 1 min. The washing step at 80% mobile phase B lasted 2 min and was followed by an equilibration step at 3% mobile phase B (starting conditions) for 6 min. The source conditions were set as follows: Gas1 = 10 psi, Gas2 = 20 psi, CurtainGas = 25 psi, SourceTemp = 100 °C, IonSpray voltage = 4500 V. The doubly charged light (613.3 m/z) and heavy precursor mass (619.3 m/z) of the SFIEDLLFNK peptide were acquired (100-1500 m/z) with a dwell time of 150 ms each.

##### B) Echo - ZenoTOF 7600

For Echo PRM analysis, a pool of the SFIEDLLFNK enriched peptide was measured six times from a 384-well plate using PRM analysis on an Echo MS+ system with the ZenoTOF 7600 mass spectrometer (Sciex, Massachusetts, USA) equipped with the Optiflow TurboV ion source operating in positive mode. AE conditions were set as following: a carrier solvent composition of 80% acetonitrile/200 nM medronic acid, a carrier solvent flow rate of 500 µL/min, a droplet count of 300 nL total ejection volume and a transducer frequency of 100 Hz. The source conditions were set as following: Gas1 = 90 psi, Gas2 = 70 psi, CurtainGas = 35 psi, CAD gas = 7 psi, SourceTemp = 300 °C, IonSpray voltage = 4500 V. The doubly charged light (613.3 m/z) and heavy precursor mass (619.3 m/z) of the SFIEDLLFNK peptide were acquired (100-1500 m/z) with a dwell time of 50 ms each.

#### DIA

##### A) SWATH

An Eksigent NanoLC 425 System (SCIEX, CA) plumbed for microflow chromatography was used and operated in trap-elute with the SCIEX TripleTOF 6600+ mass spectrometer. The samples were loaded at 10 µL/min with mobile phase A (0.1% FA in water) and trapped on a TriArt C18 guard column (YMC) (id 500 µm, length 5 mm, particle size 3 µm) for 3 min. The analytical column used was a 150 × 0.3mm Phenomenex Luna Omega Polar C18 Column (2.6 µm, 100 Å), controlled at 30°C. Peptide elution was performed at 5 µL/min using the following gradient: i) 3% to 30% mobile phase B (0.1% FA in ACN) in 60 min, ii) ramp to 80% mobile phase B in 1 min. The washing step at 80% mobile phase B lasted 4 min and was followed by an equilibration step at 3% mobile phase B (starting conditions) for 10 min. The source conditions were set as follows: GS1 = 20 psi, GS2 = 20 psi, CurtainGas = 35 psi, SourceTemp = 100 °C, IonSpray voltage = 5000 V.

For SWATH (a cycle time of 4.43 s), a 100 variable window acquisition scheme was used (see Supplementary Table 2). Briefly, SWATH MS2 spectra were collected in high sensitivity mode from 100– 1500 m/z, for 18 ms. Before each SWATH MS cycle an additional MS1 survey scan in high sensitivity mode was recorded for 100 ms.

##### B) ZenoSWATH

An ACQUITY M-Class UPLC System (Waters Corporation) was used and operated in trap-elute LC mode with the SCIEX ZenoTOF 7600 mass spectrometer. Mobile phases consisted of 0.1% formic acid in water (A) and 0.1% formic acid in acetonitrile (B). The samples were loaded at 10 µL/min with 98.5% mobile phase A and trapped on a TriArt C18 guard column (YMC) (id 500 µm, length 5 mm, particle size 3 µm) for 1.5 min. Microflow separations were done using a YMC Triart C18 (15 cm x 300 µm, 3 µm particle size) at a flow rate of 5 µL/min, using a non-linear LC gradient described in Supplementary Table 3.

An OptiFlow Turbo V ion source was used with the microflow probe (1-10 µL/min electrode) for the microflow experiments, with source parameters as following: GS1 = 20 psi, GS2 = 55 psi, Curtain Gas = 45 psi, CAD gas = 7 psi and Ionspray voltage = 4500 V. Zeno SWATH DIA experiments used 85 variable windows (Supplementary Table 4) spanning the TOF MS mass range 400-900 Da and MS/MS mass range 140-1800 Da, with Zeno trap pulsing turned on, with MS/MS accumulation times of 13 msec.

## Data processing

To visualize and analyse the MRM and PRM data, a Skyline document (Skyline Daily v.24.1.1.254) comprising both the “heavy” (from the QcontCAT protein) and “light” (from the native viral proteins) transitions of the SFIEDLLFNK, RSFIEDLLFNK, AYNVTQAFGR and ADETQALPQR peptides were compiled. Visualization of the data was achieved with PyMOL, Microsoft Excel 365 (v2311), AlphaMap (v0.1.12), and GraphPad Prism (v.10.3.1) ^44,61^.

(Zeno)SWATH data analysis was conducted using DIA-NN (v1.8.2 beta 22) in “library-free” mode. An in silico predicted spectral library was created from a compiled protein sequence database, which included both the human proteome and SwissProt-reviewed sequences of the spike protein. This generated library was subsequently employed to search and quantify the DIA data, with the “matching between runs” feature enabled to enhance sensitivity and robustness in peptide identification. The analysis parameters were set as follows: peptide lengths were specified to range from 6 to 35 amino acid residues, while the precursor m/z values were limited to 400-1200 m/z (precursor charge: 1-4+). Trypsin was selected as the digestive enzyme, allowing for a maximum of up to two missed cleavages. Fixed modification for cysteine carbamidomethylation was applied, and methionine oxidation was designated as a variable modification. The resulting DIA-NN report.tsv file from the mRNA/saRNA transfected HEK- and HeLa-cells was used for differential protein analysis using MSqRob ^48,62^. The resulting MS-DAP PDF reports, containing all analysis steps for full reproducibility and provides plenty information on how to interpret the results, and the differential abundance analysis are provided as supplementary data.

## Data availability

The mass spectrometry MRM, PRM and DIA proteomics (except the lipofectamine transfection experiments) raw data files have been deposited to the ProteomeXchange Consortium via the Panorama Public partner repository with dataset identifier PXD060927 ^63^. Data can be accessed from https://panoramaweb.org/1GQZWU.url using the reviewer account details: (Password: i*0@lSTmE1s?A6).

## Notes

The authors declare the following competing financial interest(s): Vanhulle M. and Vissers J.P.C. are employed by Waters Corporation.

## Acknowledgements

Funding from Research Foundation Flanders (FWO) [1278023N – B.V.P; G016221N – R.V.], Ghent University Special Research Fund [BOF21/DOC/076 – R.A] and European Unions’ Horizon Research and Innovation Program [OneLab - No. 101073924 and BAXERNA - No. 101080544]. We would like to thank Florian C. Sigloch from PolyQuant GmbH for providing us with the CovMS QconCAT heavy standard.

## Author Contributions

B.V.P. and M.D. contributed to conceptualization; S.C, B.S. and X.S. performed the plasmid transfection experiments, P.R. generated the 3D structure of the SPIKE protein. K.M., I.L. and R.V. performed the mRNA transfection experiments; R.A. performed the HEK293-T flow-through analysis; G.M. provided SARS-CoV-2 patient samples; P.R. provided the Pfizer-BioNTech COVID-19 Omicron XBB.1.5 mRNA vaccine. D.D. helped with funding acquisition; M.V.H. and J.P.C.V. helped with the MRM measurements. B.V.P. and M.D. wrote the original draft and all authors have proofread the manuscript.

