## Supporting Information for "Development of a General Purpose Targeted LC-MS Method for Accurate Quantification of the SARS-CoV-2 Spike Protein Expression"

Short title: SPIKE_SARS2 protein detection by LC-MS

°These authors jointly supervised the work

Keywords: Mass Spectrometry, SARS-CoV-2, LC-MS, immuno-MRM, RNA expression, SPIKE_SARS2

### Supplementary Figures

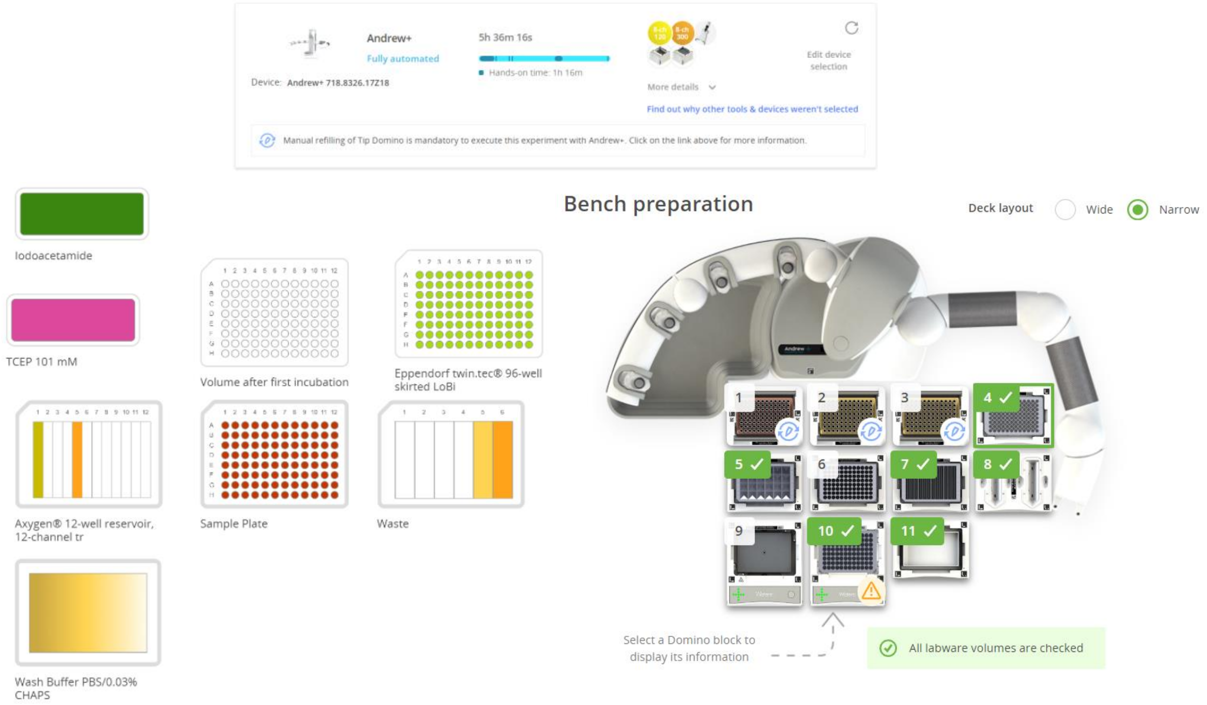


**Supplementary Figure 1. Automated SISCAPA workflow using the Andrew+™ workstation.** *(Top)* Overview of the connected devices and pipettes required to execute the Andrew+ protocol, including estimated total runtime and hands-on time. *(Left)* Schematic representation of the consumables and reagents used in each position throughout the protocol. *(Right)* Visualization of the deck layout, detailing the arrangement of modules and connected devices necessary for performing the SISCAPA SFIEDLLFNK assay.

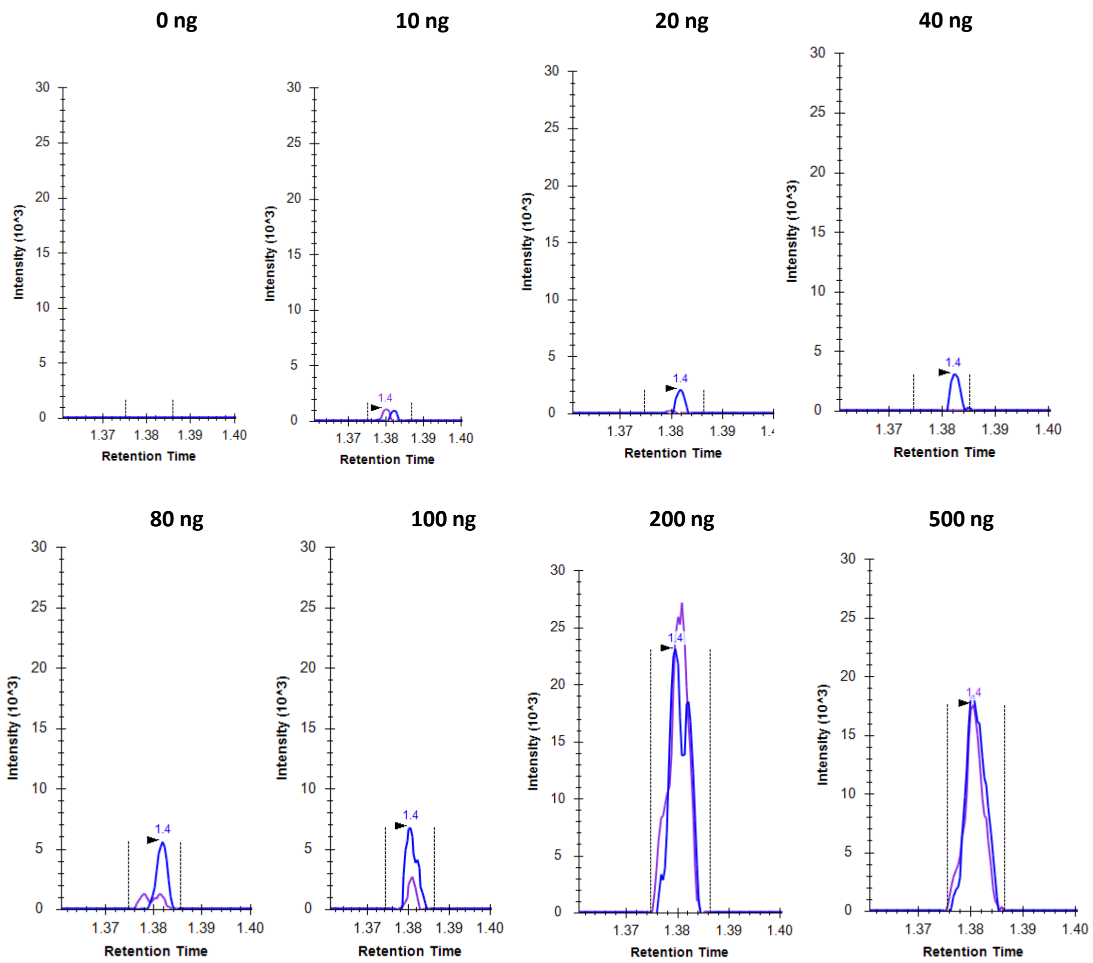


**Supplementary Figure 2. mRNA transfection dose response curve.** MRM chromatograms of peptide SFIEDLLFNK after transfecting 30,000 HEK cells with varying dosages (0-500 ng) of mRNA LNP mixture.

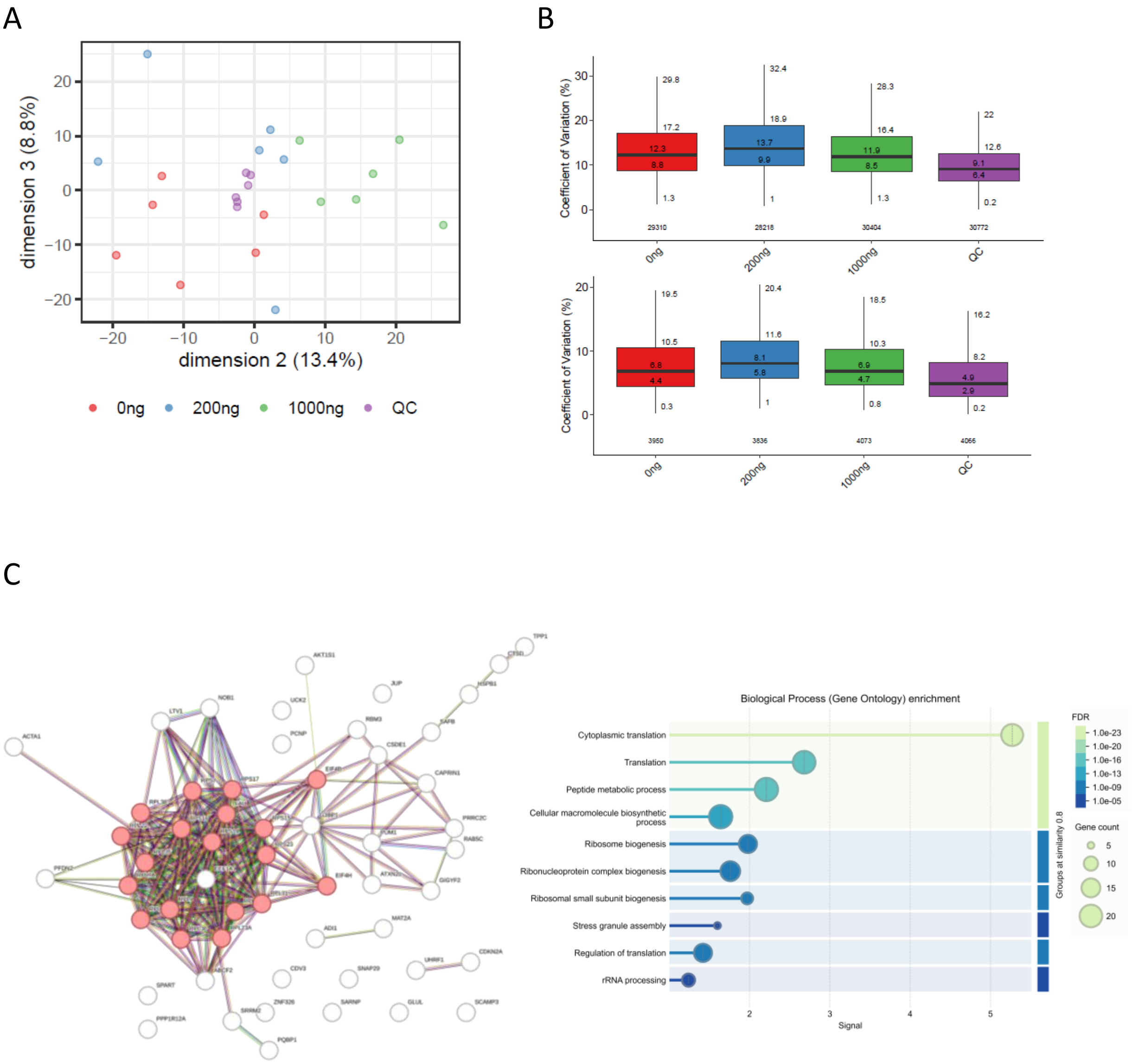


**Supplementary Figure 3. Analysis of the flow-through fractions from mRNA-transfected HEK cells following SFIEDLLFNK peptide enrichment.** A) Principal Component Analysis (PCA) of all samples, with the second and third principal components displayed on the axes, alongside their respective contribution to the explained variance. Samples cluster according to their respective conditions (0 ng, 200 ng, 1000 ng, and QC. Notably, one sample from the 200 ng condition exhibits distinct behavior, suggesting potential variation.  B) Peptide and protein coefficient of variation (%CV) analysis demonstrates robust reproducibility across all conditions, with %CV values consistently below 20%, indicating high-quality data and low variability. C) Gene Ontology (GO) enrichment analysis of all significant downregulated proteins ($q < 0.01$) in the 200 ng *vs*. 0 ng contrast, using all identified proteins as the background, reveals a significant association with cytoplasmic translation processes (red dots). This suggests a cellular response to mRNA transfection, likely reflecting translational repression. A similar trend is observed in the 1000 ng vs. 0 ng comparison (*data not shown*), further supporting the dose-dependent impact of mRNA transfection on translation-related pathways.

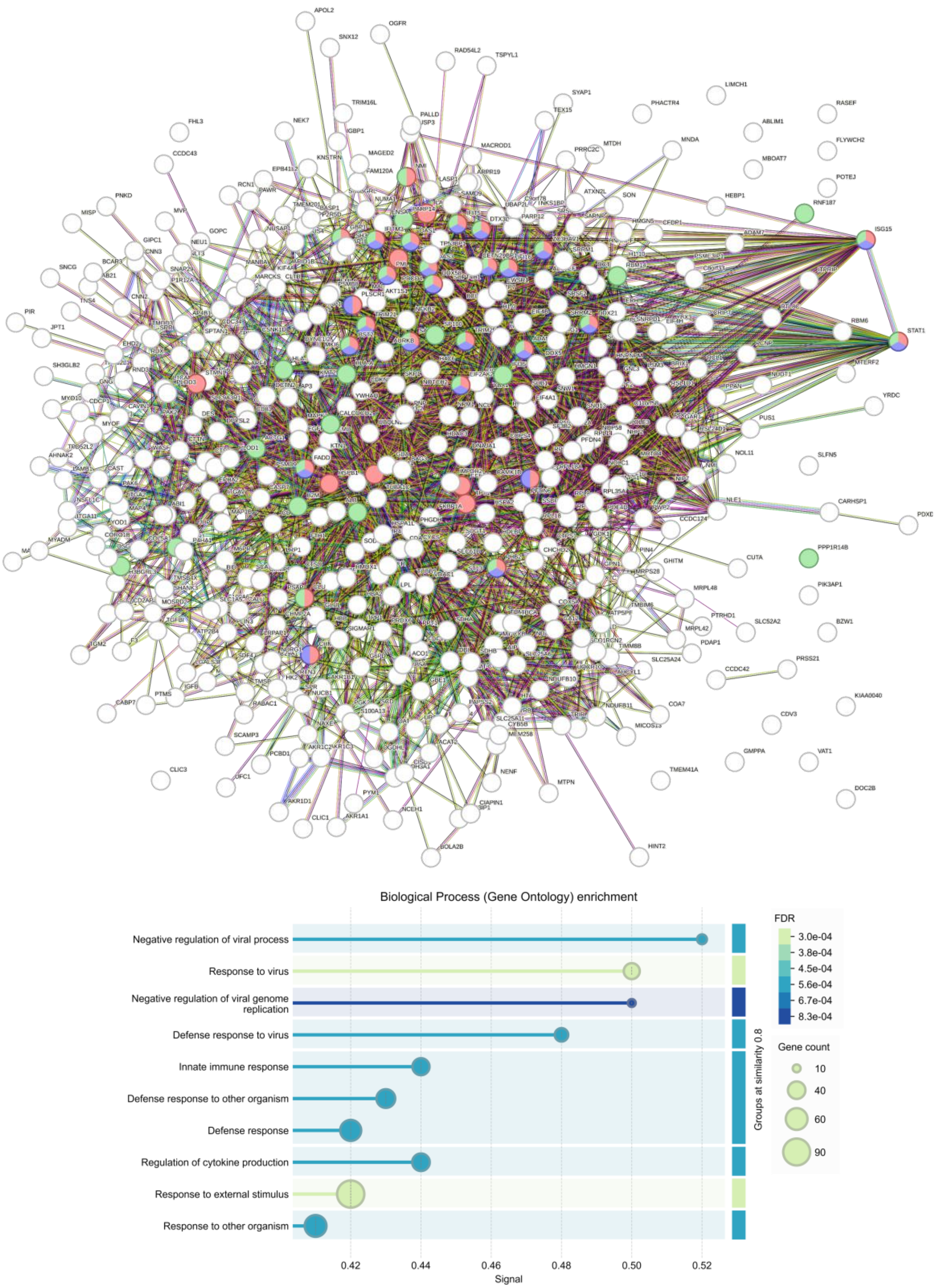


**Supplementary Figure 4. Analysis of the flow-through fractions from mRNA vs saRNA-transfected HeLa cells.** Gene Ontology (GO) enrichment analysis of all significant up- and down-regulated proteins ($q < 0.01$) in the Modified mRNA *vs*. saRNA contrast, using all identified proteins as the background, reveals a significant association with innate immune response (green dots), more specifically defending the cells against viral entry (blue and red dots), as shown by the biological process enrichment. This suggests a much more pronounced cellular response to saRNA transfection compared to Modified mRNA.

### Supplementary Tables

**Supplementary Table 1. Overview of SPIKE_SARS2 peptides detected by LC-MS.** A literature review was conducted to retrieve all proteotypic peptides detected for SPIKE_SARS2 by LC-MS to date.

| SPIKE_SARS2 | Van Puyvelde et al. | Cardozo et al. | Cazares et al. | Renuse et al. | Gouveia et al. | Zecha et al. | Saadi et al. | Pinto et al. | Long et al. | Hodgkins et al. | Schuster et al. | Pierce-Ruiz et al. | Rosen et al. | Sutton et al. | Fu et al. | Bate et al. |
|---|---|---|---|---|---|---|---|---|---|---|---|---|---|---|---|---|
| TQLPPAYTNSFTR (AA 22-34) | | | | | | | | X | | | | | | | | |
| GVYYPDK (AA 35-41) | | | | | | | X | X | | | | X | | X | | X |
| RFDNPVLPFNDGVYFASTEK (AA 78-97) | | | | X | | | X | | | | | | | | | |
| GWIFGTTLDSK (AA 103-113) | X | | | X | X | | X | | | | | | | | | X |
| VCEFQFCNDPFLGVYYHK (AA 130-147) | | | | | | | X | | | | | | | | | |
| SWMESEFR (AA 151-158) | | | | | | | X | | | | | | | | | |
| NIDGYFK (AA 195-202) | | | X | | | | X | | | | | | | | | |
| HTPINLVR (AA 207-214) | | | | | X | | X | X | | | X | | X | | | |
| FQTLLALHR (AA 238-246) | | | X | X | X | | X | X | | | X | | | | | |
| SYLTPGDSSSGWTAGA AAYYVGYLQPR (AA 247-273) | | | | | | | X | | | | | | | | | |
| TFLLK (AA 274-278) | | | | | | | | X | | | | | | | | |
| SFTVEK (AA 305-310) | | | | | | | X | | | | | | | | | |
| GIYQTSNFR (AA 311-319) | | | | | | X | X | X | | | | X | | X | | |
| VQPTESIVR (AA 320-328) | | | | | | X | X | | | | | | | | | |
| FASVYAWNR (AA 347-355) | | | | X | | X | X | X | | | | | | | | |
| CYGVSPTK (AA 379-386) | | | | | | | X | X | | | | | | | | |
| LNDLCFTNVYADSFVIR (AA 387-403) | | | | | | | X | | | | | | | | | |
| IADYNYK (AA 418-424) | | | | | | | X | X | | | | | | | | |
| QIAPGQTGK (AA 409-415) | | | | | | | | | X | | | | | | | |
| LPDDFTGCVIAWNSNNLDSK (AA 425-444) | | | | | | | X | | | | | | | | | |
| VGGNYNYLYR (AA 445-454) | | X | | | | X | X | X | | | | | | | | |
| SNLKPFER (AA 459-466) | | | X | | | | X | | | | | | | | | |
| VVVLSFELLHAPATVCGPK (AA 510-528) | | | | | | X | X | | | | | | | | | |
| STNLVKNK (AA 530-537) | | | | | | | | X | | | | | | | | |
| FLPFQQFGR (AA 559-567) | | | | X | | X | X | X | X | X | X | X | X | X | X | |
| DIADTTDAVR (AA 568-577) | | | X | | | | X | | | | | | | | | |
| VYSTGSNVFQTR (AA 635-646) | | X | | X | | X | X | X | | | | | | | | |
| ALTGIAVEQDKNTQEVFAQVK (AA 766-786) | | | | X | | | X | | | | | | | | | |
| SFIEDLLFNK (AA 816-825) | X | | | | | | X | | | | X | X | X | X | | X |
| VTLADAGFIK (AA 826-835) | | | | | | | | | | | | X | | X | X | |
| QYGDCLGDIAAR (AA 836-847) | | | | | | | X | | | | | | | | | |
| DLICAQK (AA 848-854) | | | | | | | | | | | | | | | | |
| FNGIGVTQNVLYENQK (AA 906-921) | | X | | | | | X | | | | | | | | | |
| LIANQFNSAIGK (AA 922-933) | | | | | | | X | X | | | | | | | | |
| LQDVVNQNAQALNTLVK (AA 948-964) | | X | | X | | | | | | | | | | | | |
| LQSLQTYVTQQLIR (AA 1001-1014) | | X | | X | X | | X | X | | | | | | | | |
| ASANLAATK (AA 1020-1028) | | | | | | | | X | | | | X | X | X | | X |
| MSECVLGQSK (AA 1029-1038) | | | | | | | X | X | | | | | | | | |
| RVDFCGK (AA 1039-1045) | | | | | | | | X | | | | | | | | |
| VDFCGK (AA 1040-1045) | | | | | | | | X | | | | | | | | |
| GYHLMSFPQSAPHGVVFLHVTYVPAQEK (AA 1046-1073) | | | | | | | X | | | | | | | | | |
| EIDRLNEVAK (AA 1182-1191) | | | | | | | | X | | | | | | | | |
| LNEVAK (AA 1186-1191) | | | | | | | | X | | | | | | | | |
| FDEDDSEPVLK (AA 1256-1266) | | | | | | | X | | | | | | | | | |

**Supplementary Table 2. 100-Variable Window (VW) scheme used for SWATH analysis of the flow-through of plasmid transfected HEK cells.**

| | Start Mass (Da) | Stop Mass (Da) | CES |
|---|---|---|---|
| Exp 1 | 399.5 | 406.5 | 0 |
| Exp 2 | 405.5 | 412.5 | 0 |
| Exp 3 | 411.5 | 418.5 | 0 |
| Exp 4 | 417.5 | 424.5 | 0 |
| Exp 5 | 423.5 | 430.5 | 0 |
| Exp 6 | 429.5 | 436.5 | 0 |
| Exp 7 | 435.5 | 442.5 | 0 |
| Exp 8 | 441.5 | 448.5 | 0 |
| Exp 9 | 447.5 | 454.5 | 0 |
| Exp 10 | 453.5 | 459.5 | 0 |
| Exp 11 | 458.5 | 464.5 | 0 |
| Exp 12 | 463.5 | 469.5 | 0 |
| Exp 13 | 468.5 | 474.5 | 0 |
| Exp 14 | 473.5 | 479.5 | 0 |
| Exp 15 | 478.5 | 484.5 | 0 |
| Exp 16 | 483.5 | 489.5 | 0 |
| Exp 17 | 488.5 | 494.5 | 0 |
| Exp 18 | 493.5 | 499.5 | 0 |
| Exp 19 | 498.5 | 504.5 | 0 |
| Exp 20 | 503.5 | 509.5 | 0 |
| Exp 21 | 508.5 | 514.5 | 0 |
| Exp 22 | 513.5 | 519.5 | 0 |
| Exp 23 | 518.5 | 524.5 | 0 |
| Exp 24 | 523.5 | 529.5 | 0 |
| Exp 25 | 528.5 | 534.5 | 0 |
| Exp 26 | 533.5 | 539.5 | 0 |
| Exp 27 | 538.5 | 544.5 | 0 |
| Exp 28 | 543.5 | 549.5 | 0 |
| Exp 29 | 548.5 | 554.5 | 0 |
| Exp 30 | 553.5 | 559.5 | 0 |
| Exp 31 | 558.5 | 564.5 | 0 |
| Exp 32 | 563.5 | 569.5 | 0 |
| Exp 33 | 568.5 | 574.5 | 0 |
| Exp 34 | 573.5 | 579.5 | 0 |
| Exp 35 | 578.5 | 584.5 | 0 |
| Exp 36 | 583.5 | 589.5 | 0 |
| Exp 37 | 588.5 | 594.5 | 0 |
| Exp 38 | 593.5 | 599.5 | 0 |

| | | | |
|---|---|---|---|
| Exp 39 | 598.5 | 604.5 | 0 |
| Exp 40 | 603.5 | 609.5 | 0 |
| Exp 41 | 608.5 | 614.5 | 0 |
| Exp 42 | 613.5 | 619.5 | 0 |
| Exp 43 | 618.5 | 624.5 | 0 |
| Exp 44 | 623.5 | 629.5 | 0 |
| Exp 45 | 628.5 | 634.5 | 0 |
| Exp 46 | 633.5 | 639.5 | 0 |
| Exp 47 | 638.5 | 644.5 | 0 |
| Exp 48 | 643.5 | 649.5 | 0 |
| Exp 49 | 648.5 | 654.5 | 0 |
| Exp 50 | 653.5 | 660.5 | 0 |
| Exp 51 | 659.5 | 666.5 | 0 |
| Exp 52 | 665.5 | 672.5 | 0 |
| Exp 53 | 671.5 | 678.5 | 0 |
| Exp 54 | 677.5 | 684.5 | 0 |
| Exp 55 | 683.5 | 690.5 | 0 |
| Exp 56 | 689.5 | 696.5 | 0 |
| Exp 57 | 695.5 | 702.5 | 0 |
| Exp 58 | 701.5 | 708.5 | 0 |
| Exp 59 | 707.5 | 714.5 | 0 |
| Exp 60 | 713.5 | 720.5 | 0 |
| Exp 61 | 719.5 | 726.5 | 0 |
| Exp 62 | 725.5 | 732.5 | 0 |
| Exp 63 | 731.5 | 738.5 | 0 |
| Exp 64 | 737.5 | 744.5 | 0 |
| Exp 65 | 743.5 | 750.5 | 0 |
| Exp 66 | 749.5 | 756.5 | 0 |
| Exp 67 | 755.5 | 763.5 | 0 |
| Exp 68 | 762.5 | 770.5 | 0 |
| Exp 69 | 769.5 | 777.5 | 0 |
| Exp 70 | 776.5 | 784.5 | 0 |
| Exp 71 | 783.5 | 791.5 | 0 |
| Exp 72 | 790.5 | 798.5 | 0 |
| Exp 73 | 797.5 | 805.5 | 0 |
| Exp 74 | 804.5 | 812.5 | 0 |
| Exp 75 | 811.5 | 819.5 | 0 |
| Exp 76 | 818.5 | 826.5 | 0 |
| Exp 77 | 825.5 | 834.5 | 0 |
| Exp 78 | 833.5 | 842.5 | 0 |
| Exp 79 | 841.5 | 850.5 | 0 |

| | | | |
|---|---|---|---|
| Exp 80 | 849.5 | 858.5 | 0 |
| Exp 81 | 857.5 | 867.5 | 0 |
| Exp 82 | 866.5 | 876.5 | 0 |
| Exp 83 | 875.5 | 885.5 | 0 |
| Exp 84 | 884.5 | 894.5 | 0 |
| Exp 85 | 893.5 | 903.5 | 0 |
| Exp 86 | 902.5 | 914.5 | 0 |
| Exp 87 | 913.5 | 925.5 | 0 |
| Exp 88 | 924.5 | 936.5 | 0 |
| Exp 89 | 935.5 | 950.5 | 0 |
| Exp 90 | 949.5 | 964.5 | 0 |
| Exp 91 | 963.5 | 978.5 | 0 |
| Exp 92 | 977.5 | 992.5 | 0 |
| Exp 93 | 991.5 | 1011.5 | 0 |
| Exp 94 | 1010.5 | 1030.5 | 0 |
| Exp 95 | 1029.5 | 1054.5 | 0 |
| Exp 96 | 1053.5 | 1078.5 | 0 |
| Exp 97 | 1077.5 | 1117.5 | 0 |
| Exp 98 | 1116.5 | 1156.5 | 0 |
| Exp 99 | 1155.5 | 1200.5 | 0 |
| Exp 100 | 1199.5 | 1249.5 | 0 |

**Supplementary Table 3. LC gradient used for ZenoSWATH analysis of the flow-through.**

| Time (min) | %A | %B |
|---|---|---|
| **0** | 98.5 | 1.5 |
| **0.5** | 92.7 | 7.3 |
| **1** | 89.5 | 10.5 |
| **2** | 87 | 13 |
| **3** | 85.3 | 14.7 |
| **5** | 83.2 | 16.8 |
| **7.5** | 80.9 | 19.1 |
| **10** | 78.9 | 21.1 |
| **12.5** | 76.5 | 23.5 |
| **15** | 73.8 | 26.2 |
| **17** | 70 | 30 |
| **18** | 66.7 | 33.3 |
| **19** | 62.6 | 37.4 |
| **19.5** | 60 | 45 |
| **20** | 55 | 42 |
| **21** | 10 | 90 |
| **25** | 10 | 90 |
| **26** | 98.5 | 1.5 |
| **30** | 98.5 | 1.5 |

**Supplementary Table 4. 85-Variable Window (VW) scheme used for the Zeno SWATH analysis of the flow-through of mRNA transfected HEK cells.**

| | Start Mass (Da) | Stop Mass (Da) | CES |
|---|---|---|---|
| Exp 1 | 399.5 | 406.5 | 0 |
| Exp 2 | 405.5 | 412.5 | 0 |
| Exp 3 | 411.5 | 418.5 | 0 |
| Exp 4 | 417.5 | 424.5 | 0 |
| Exp 5 | 423.5 | 430.5 | 0 |
| Exp 6 | 429.5 | 436.5 | 0 |
| Exp 7 | 435.5 | 442.5 | 0 |
| Exp 8 | 441.5 | 448.5 | 0 |
| Exp 9 | 447.5 | 454.5 | 0 |
| Exp 10 | 453.5 | 459.5 | 0 |
| Exp 11 | 458.5 | 464.5 | 0 |
| Exp 12 | 463.5 | 469.5 | 0 |
| Exp 13 | 468.5 | 474.5 | 0 |
| Exp 14 | 473.5 | 479.5 | 0 |
| Exp 15 | 478.5 | 484.5 | 0 |
| Exp 16 | 483.5 | 489.5 | 0 |
| Exp 17 | 488.5 | 494.5 | 0 |
| Exp 18 | 493.5 | 499.5 | 0 |
| Exp 19 | 498.5 | 504.5 | 0 |
| Exp 20 | 503.5 | 509.5 | 0 |
| Exp 21 | 508.5 | 514.5 | 0 |
| Exp 22 | 513.5 | 519.5 | 0 |
| Exp 23 | 518.5 | 524.5 | 0 |
| Exp 24 | 523.5 | 529.5 | 0 |
| Exp 25 | 528.5 | 534.5 | 0 |
| Exp 26 | 533.5 | 539.5 | 0 |
| Exp 27 | 538.5 | 544.5 | 0 |
| Exp 28 | 543.5 | 549.5 | 0 |
| Exp 29 | 548.5 | 554.5 | 0 |
| Exp 30 | 553.5 | 559.5 | 0 |
| Exp 31 | 558.5 | 564.5 | 0 |
| Exp 32 | 563.5 | 569.5 | 0 |
| Exp 33 | 568.5 | 574.5 | 0 |
| Exp 34 | 573.5 | 579.5 | 0 |
| Exp 35 | 578.5 | 584.5 | 0 |
| Exp 36 | 583.5 | 589.5 | 0 |
| Exp 37 | 588.5 | 594.5 | 0 |
| Exp 38 | 593.5 | 599.5 | 0 |

| | | | |
|---|---|---|---|
| Exp 39 | 598.5 | 604.5 | 0 |
| Exp 40 | 603.5 | 609.5 | 0 |
| Exp 41 | 608.5 | 614.5 | 0 |
| Exp 42 | 613.5 | 619.5 | 0 |
| Exp 43 | 618.5 | 624.5 | 0 |
| Exp 44 | 623.5 | 629.5 | 0 |
| Exp 45 | 628.5 | 634.5 | 0 |
| Exp 46 | 633.5 | 639.5 | 0 |
| Exp 47 | 638.5 | 644.5 | 0 |
| Exp 48 | 643.5 | 649.5 | 0 |
| Exp 49 | 648.5 | 654.5 | 0 |
| Exp 50 | 653.5 | 660.5 | 0 |
| Exp 51 | 659.5 | 666.5 | 0 |
| Exp 52 | 665.5 | 672.5 | 0 |
| Exp 53 | 671.5 | 678.5 | 0 |
| Exp 54 | 677.5 | 684.5 | 0 |
| Exp 55 | 683.5 | 690.5 | 0 |
| Exp 56 | 689.5 | 696.5 | 0 |
| Exp 57 | 695.5 | 702.5 | 0 |
| Exp 58 | 701.5 | 708.5 | 0 |
| Exp 59 | 707.5 | 714.5 | 0 |
| Exp 60 | 713.5 | 720.5 | 0 |
| Exp 61 | 719.5 | 726.5 | 0 |
| Exp 62 | 725.5 | 732.5 | 0 |
| Exp 63 | 731.5 | 738.5 | 0 |
| Exp 64 | 737.5 | 744.5 | 0 |
| Exp 65 | 743.5 | 750.5 | 0 |
| Exp 66 | 749.5 | 756.5 | 0 |
| Exp 67 | 755.5 | 763.5 | 0 |
| Exp 68 | 762.5 | 770.5 | 0 |
| Exp 69 | 769.5 | 777.5 | 0 |
| Exp 70 | 776.5 | 784.5 | 0 |
| Exp 71 | 783.5 | 791.5 | 0 |
| Exp 72 | 790.5 | 798.5 | 0 |
| Exp 73 | 797.5 | 805.5 | 0 |
| Exp 74 | 804.5 | 812.5 | 0 |
| Exp 75 | 811.5 | 819.5 | 0 |
| Exp 76 | 818.5 | 826.5 | 0 |
| Exp 77 | 825.5 | 834.5 | 0 |
| Exp 78 | 833.5 | 842.5 | 0 |
| Exp 79 | 841.5 | 850.5 | 0 |

| | | | |
|---|---|---|---|
| Exp 80 | 849.5 | 858.5 | 0 |
| Exp 81 | 857.5 | 867.5 | 0 |
| Exp 82 | 866.5 | 876.5 | 0 |
| Exp 83 | 875.5 | 885.5 | 0 |
| Exp 84 | 884.5 | 894.5 | 0 |
| Exp 85 | 893.5 | 903.5 | 0 |
